# Adaptation and latitudinal gradients in species interactions: nest predation in birds

**DOI:** 10.1101/552380

**Authors:** Benjamin G. Freeman, Micah N. Scholer, Mannfred M. A. Boehm, Julian Heavyside, Dolph Schluter

## Abstract

The “biotic interactions” hypothesis proposes that strong species interactions in the tropics drive faster divergence in the tropics. However, support for the idea that interactions are stronger in the tropics is mixed. Here we propose an explanation for why observed interaction strengths might often be similar across latitudes. We suggest populations might adapt to latitudinal differences in species interaction regimes, which can have the effect of flattening observed latitudinal gradients in interaction rates. To investigate this idea, we examine a canonical example of a strong tropical biotic interaction—nest predation rates in land birds. Surprisingly, we find that daily rates of nest predation vary minimally with latitude. We then consider the possibility that evolved life history differences across latitudes contribute to the flat gradient. We focus on the duration of the nesting period, which is longer in the tropics than in the temperate zone and thought to represent a component of adaptation to tropical nest predators. Greater nesting period duration is known to be associated with lower daily predation rates. When we statistically control for nesting period duration, we recover a pattern where daily rates of nest predation are highest in the tropics. The implication of this analysis is that the evolution of longer nesting periods in tropical species (and shorter nesting periods in the temperate zone) has helped to flatten a "baseline" latitudinal gradient in daily rates of nest predation. More generally, we propose that adaptation to latitudinal differences in biotic interactions can flatten latitudinal gradients in interaction strength.

---

Since Darwin, ecologists have suggested that biotic interactions increase in strength towards the Equator (Darwin 1859; Wallace 1869; Dobzhansky 1950; MacArthur 1972). Dobzhansky (1950) expressed this viewpoint by arguing “Where physical conditions are easy, interrelationships between species become the paramount adaptive problem…This is probably the case in most tropical communities.” Strong biotic interactions in the tropics are hypothesized to generate strong selection that, in turn, leads to faster rates of evolution and speciation in the tropics (Schemske 2009). If so, strong biotic interactions in the tropics may explain in part why there are far more species at low latitudes than in the temperate zone (Schemske 2009).

The biotic interactions hypothesis has inspired a growing number of studies that test the prediction that total biotic interactions are indeed strongest in the tropics (Schemske et al. 2009; Moles and Ollerton 2016). Two principal approaches use measurements of the rates of biotic interactions to assess interaction strength across latitudes. The first is to place a standardized artificial model at many sites, such as an artificial prey type, and measure the frequency of biotic interactions, such as predator attack rates, experienced by this model (Roslin et al. 2017; Hargreaves et al. 2019). The second is to measure the frequency of biotic interactions that wild populations actually experience, repeating this at a large number of sites across latitudes to account for variability (e.g., Kubelka et al. 2018). A fundamental difference between these two approaches is that standardized models are evolutionarily naïve and measure rates of biotic interaction experienced by populations that have not adapted to local interactions. By contrast, wild studies can measure rates of biotic interaction experienced by populations that have had the opportunity to adapt to local species interactions. The possibility exists that adaptation in response to interacting species could reduce, or perhaps even eliminate, any latitudinal gradient in interaction rates. This might explain why several studies using naïve models report higher interaction rates at low latitudes (e.g., Roslin et al. 2017; Hargreaves et al. 2019) whereas many observational studies of wild populations report latitudinal gradients in interaction rates that are more flat (reviewed by Moles and Ollerton 2016).

Here we study rates of nest predation experienced by land birds, which are widely thought to be greatest at low latitudes (Skutch 1985; Robinson et al. 2000; Schemske et al. 2009; Mckinnon et al. 2010; Remeš et al. 2012; DeGregorio et al. 2016; Kubelka et al. 2018; but see Martin et al. 2017). The drivers of purported high nest predation in the tropics are not certain, but one possible cause is the apparently higher diversity of nest predators in the tropics, which include tropical predator species that appear to prey almost exclusively upon eggs (DeGregorio et al. 2016; Menezes and Marini 2017). To test these ideas, we compiled observational data on daily rates of nest survival in wild Western Hemisphere birds from the published literature. We use daily rates of nest survival to estimate daily rates of nest predation, as the bulk of nest failure is due to predation and this is consistent across latitudes (see methods). Tropical birds must deal with a distinct and diverse community of nest predators, and we explore whether tropical birds have evolved life history traits that reduce the realized rates of nest predation they experience in nature. We then statistically control for one divergent life history trait to estimate a “baseline” latitudinal gradient in rates of nest predation, and compare this to the observed (“derived”) latitudinal gradient in rates of nest predation. Our synthesis thus investigates not only latitudinal patterns in the intensity of a biotic interaction, but also the interplay between ecological interaction and evolutionary consequence across latitudinal gradients.

## Methods

### Assembling nest predation data

We searched the peer-reviewed literature to find studies that have measured nest predation for land bird populations breeding in continental North, Central or South America. We focus on the Western Hemisphere because nest predation data for tropical birds outside of the Americas (e.g., from the Asian and African tropics) is scarce (for data from Australia see Remeš et al. 2012). We included only studies of real nests because we were interested in biotic interactions experienced by populations in nature and because predation on artificial nests is poorly correlated with predation on real nests (King et al. 1999; Burke et al. 2004; Moore and Robinson 2004; Robinson et al. 2005). Following previous studies investigating latitudinal trends in nest predation, we restricted our analysis to land birds that are not cavity nesters—primarily passerines, but also a small number of doves, hummingbirds, and other non-passerines (Robinson et al. 2000; Remeš et al. 2012; Martin et al. 2017). Several previous studies have analyzed nest predation data from the Americas (Nice 1957; Skutch 1985; Kulesza 1990; Conway and Martin 2000; Robinson et al. 2000; Boyle et al. 2016; Martin et al. 2017). These syntheses are valuable summaries of relevant studies, but did not always present the complete set of data that we were interested in for each study. Hence, we extracted data from the original publications in all cases. We only included studies that reported data based on at least 10 nests. When studies found that nest predation differed between “natural” and “disturbed” habitats (typically differently sized forest fragments), we used data from the “natural” habitat. Last, we did not include data from studies in which authors explicitly stated that nest failure was due to human activities such as mowing (e.g., birds nesting in hay fields), or, in one case, when “much of the predation is known to have been by young boys” (Peterson and Young 1950).

We located additional studies by conducting a Web of Science search in February 2018 with the keywords “Nest predation” OR “Breeding ecology” OR “Nest success” AND “bird”. Because there is a latitudinal gradient in data availability (more data in the temperate zone, less in the tropics), we expended additional effort to more exhaustively search for tropical studies. Specifically, we: (1) conducted additional country-specific Web of Science searches for each nation in Central and South America, with keywords “Nest predation” OR “Nest success” AND “Country Name”, where “Country Name” was the name of each individual Central and South American country; (2) examined the entire publication records of scientists who have extensively studied Neotropical bird breeding biology; and (3) followed citation webs to search for additional relevant studies from the tropics and southern temperate zone.

For each species from each study that met our criteria described above, we extracted the following information: (1) Species name; (2) Sample size of nests; (3) Nest success, presented either as fledging success (often termed “apparent success”, an estimate of the percentage of nests that successfully fledge young) or as daily survival rate (the probability that an egg or nestling survives from one day to the next); (4) Latitude and longitude; and (5) Clutch size, incubation and nestling periods, extracted from the paper itself or from Handbook of the Birds of the World Alive (del Hoyo et al. 2018).

Our final dataset included nest predation data for 515 unique species-site combinations (from 244 studies and representing 314 species). The bulk of this dataset comes from studies conducted between ~40**°** South and ~50**°** North. The majority of studies came from the Northern Hemisphere temperate zone (269 unique species-site combinations), but the tropics (absolute latitude < 23.4; 187 unique species-site combinations) and Southern Hemisphere temperate zone (59 unique species-site combinations) were also well represented (Fig 1). This dataset included 369 estimates of fledging success and 267 estimates of daily survival rate (see Fig S1).

**Figure 1.**
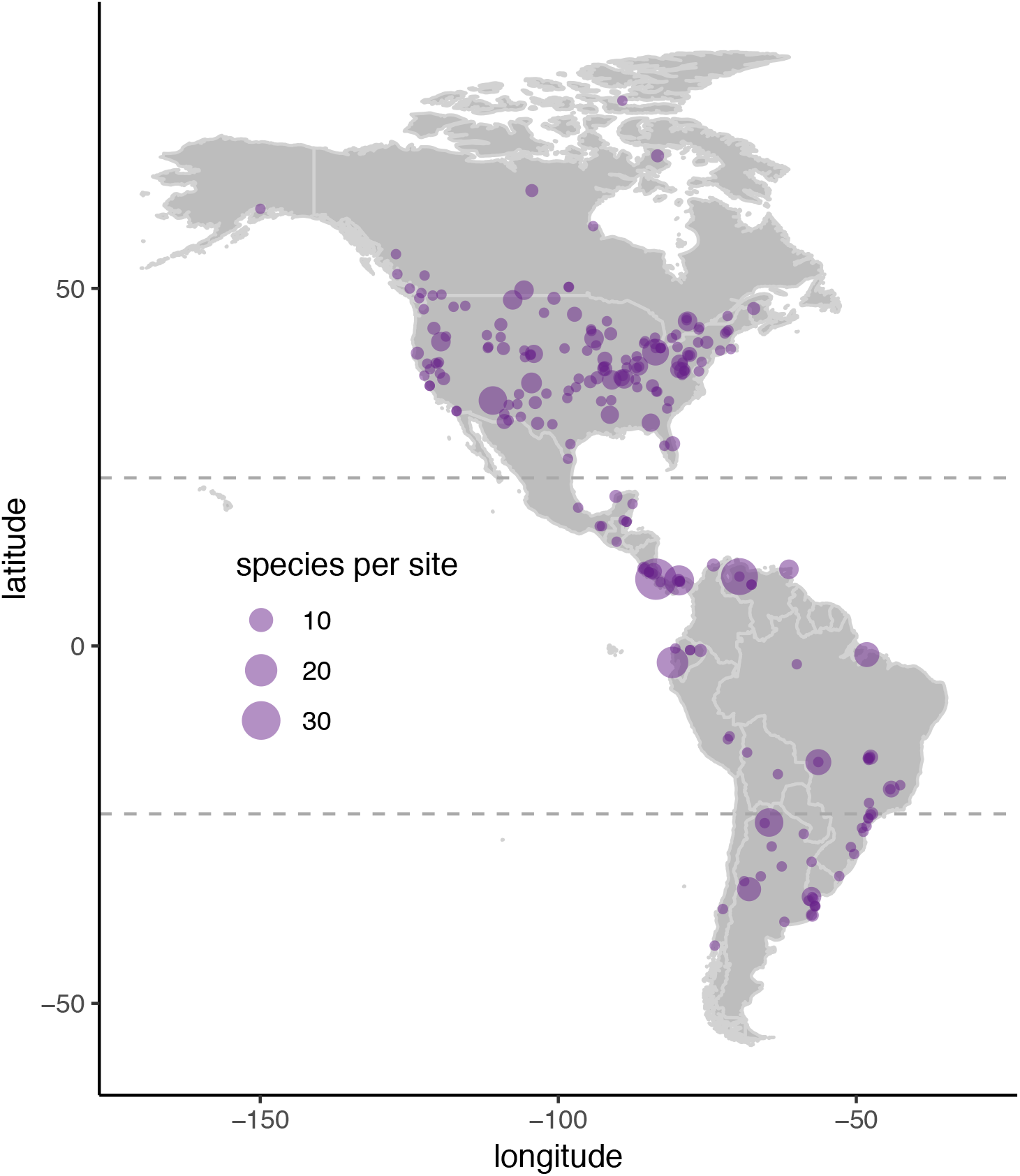
Map of studies measuring nest predation experienced by land birds in the Americas (dataset = 244 studies). Many studies report data for multiple species from the same site, illustrated by the size of the circle. The Tropics of Cancer and Capricorn (at 23.4**°** N and S, respectively) delimit the tropics, and are illustrated with dashed lines.

### Nest predation vs. nest failure

Nest predation is difficult to study directly, as it is difficult to identify the cause of nest failure. Predators are responsible for the large majority of nest mortality in most places (Remeš et al. 2012; Martin et al. 2017), In our dataset, around one quarter of studies (66 out of 244) attempted to identify causes of nest failure. Most nest failure was attributed to predation (73%; *N* = 112 unique species-site combinations), a figure remarkably similar to that previously reported for Australian birds (72%; Remeš et al. 2012). Further, the percentage of nest failures due to predation was similar across latitudes (slope estimate for latitude in a univariate linear model with percentage of nest failure attributed to predation as the response variable and latitude as a predictor variable = −0.00083 ± 0.00074, *p* = 0.27). Hence, we follow previous analyses and classify all nest failures as due to predation (Martin et al. 2017). This approach likely slightly overestimates nest predation rates. However, it does not appear to introduce any bias, and offers the advantage of greater clarity, as our focus in this paper is on predators and nest predation.

### Latitudinal variation in daily rates of nest predation

We assessed whether daily rates of nest predation (i.e., predation of the bird egg or nestling) vary as a function of latitude by fitting mixed effect meta-analytic models using the “metafor” package (Viechtbauer 2010) in R (R Development Core Team 2017). Data for this analysis were 267 estimates of daily survival rate, which we used to calculate daily predation rate (132 from the north temperate zone, 90 from the tropics, and 45 from the south temperate zone). The meta-analytic models weight individual estimates by the inverse of their squared standard errors, and incorporate the estimated variance among the study-specific effect sizes. We fit three models that correspond to different biological hypotheses: (1) no latitudinal gradient in predation (intercept-only model); (2) a linear latitudinal gradient in predation (single slope term, equal slopes in both Northern and Southern hemispheres); and (3) a breakpoint linear model wherein predation differs between the tropics and temperate zone (zero slope within the tropics, ≤ 23.4 degrees absolute latitude, equal slopes for temperate latitudes in both Northern and Southern hemispheres). We then compared model fits using AIC. After determining the best-fit model, we fit an expanded version of the best-fit model to test the influence of non-independence among effect sizes. This model included species as a random effect, and we incorporated phylogenetic relationships between species by specifying shared phylogenetic branch length in the variance-covariance matrix. Branch lengths were measured from a majority rules consensus tree calculated from 1000 phylogenies with the “Hackett” backbone pruned to our study taxa and downloaded from birdtree.org (Jetz et al. 2012). We did not include phylogenetic relationships in any of our initial models of latitudinal variation in interaction rate, because we do not believe our question requires a phylogenetic approach (e.g., in principle we could test for latitudinal variation in rates of nest predation even if there was only one species with a broad latitudinal distribution in the dataset). We then incorporated phylogenetic relationships into secondary models to probe one potential explanation of our results, that latitudinal differences in nest predation are related to phylogenetic differences between tropical and temperate taxa.

### Latitudinal variation in nesting period duration

We found minimal variation in daily predation rates among latitudes (see Results). To explore causes of this surprising result, we examined latitudinal patterns in nesting period duration. We calculated nesting period length as the number of days from the laying of the first egg until fledging, assuming that species begin incubating upon laying the final egg. We first plotted species’ nesting period durations vs. latitude (using data on nesting period for 292 species from our dataset for which we could calculated nesting period; 110 from the north temperate zone, 142 from the tropics, and 40 from the south temperate zone), and observed clear latitudinal patterns in nesting period duration, with longer nesting periods in the tropics (see Results). We therefore tested the evolutionary association between latitude and nesting period duration by fitting a phylogenetic generalized least squares (PGLS) regression using the “ape” package (Paradis et al. 2004). The response variable in this model was nesting period duration, absolute value of latitude was a fixed effect, Pagel’s λ was estimated using maximum likelihood, and we used a majority rules consensus tree from 1000 phylogenies with the “Hackett” backbone downloaded from birdtree.org (Jetz et al. 2012). In both reality and our dataset, oscine passerines predominate in the temperate zone whereas suboscines are more common in the tropics. To explore whether our overall results are mirrored within these groups, we repeated the PGLS for oscines only (*N* = 175 species) and for suboscines only (*N* = 86 species).

To quantify the relationship between nesting period duration and daily predation rates, we fit a meta-analytic model to predict daily predation rate using the “metafor” package. Data for this analysis came from 175 species for which we had both estimates of daily survival rate (used to calculate daily predation rate) and information on nesting period length (132 from the north temperate zone, 84 from the tropics, and 36 from the south temperate zone). We first fit a model that estimated different slopes and intercepts for different latitudinal zones (predictor variables = nesting period duration, latitudinal zone [north temperate, tropical, south temperate], and an interaction between nesting period duration and latitudinal zone). We next fit a model where latitudinal zones had different intercepts but the same slope (i.e., without the interaction term between nesting period duration and latitudinal zone), and compared fit of the “different slopes” and “same slopes” models using the “anova” function in R. Last, we fit an expanded version of the “same slopes” model that included species as a random effect and incorporated phylogeny by specifying phylogenetic branch length as the variance-covariance matrix. Again, branch lengths were measured from a majority rules consensus tree calculated from 1000 phylogenies with the “Hackett” backbone”, pruned to our study taxa and downloaded from birdtree.org (Jetz et al. 2012)

### Latitudinal variation in fledging success

We examined latitudinal patterns in fledging success by repeating the three meta-analytic models described above for latitudinal patterns in daily predation rate, but with fledging success as the response variable. Data for this analysis were 369 estimates of fledging success (187 from the north temperate zone, 152 from the tropics, and 30 from the south temperate zone). Upon plotting the data for the best-fit model (breakpoint regression with equal slopes in Northern and Southern temperate zones), we noticed that this model provided a poor fit to data from the Southern temperate zone. We therefore fit an additional breakpoint regression model where we allowed slopes to differ between Northern and Southern temperate zones, and compared model fits using AIC. Last, we fit an expanded version of the best-fit model that included species as a random effect and specified phylogenetic branch lengths as the variance-covariance matrix (calculated as previously described).

## Results

### Latitudinal variation in daily rates of nest predation is minimal

We found evidence that daily rates of nest predation in land birds are similar across latitudes within the Western Hemisphere. The most strongly supported model of daily nest survival fit a line with equal daily predation rate (~ 0.04) across the entire latitudinal gradient (Fig 2, Table 1). Alternative models that fit symmetric, non-zero slopes to the relationship between latitude and daily survival rates were less well supported (ΔAIC ~ 7; Table 1). These less supported models were similar to the best-fit model in that they estimated slopes that were nearly flat (Table S1), with estimated daily nest predation at 45**°** North only slightly lower (by ~ 0.004) than at the equator. Our results did not change when including phylogenetic relationships and species identity in our model (Table S2).

**Table 1.** Model comparison of metafor models with daily predation rate as the response variable. The best-fit model fit a constant value of daily predation rate across the entire latitudinal expanse of the dataset.

| Model | $\Delta AIC$ |
| --- | --- |
| Constant (daily predation rates do not vary with latitude) | -- |
| Linear regression | 6.89 |
| Breakpoint regression (breakpoint = 23.4°) | 7.54 |

**Figure 2.**
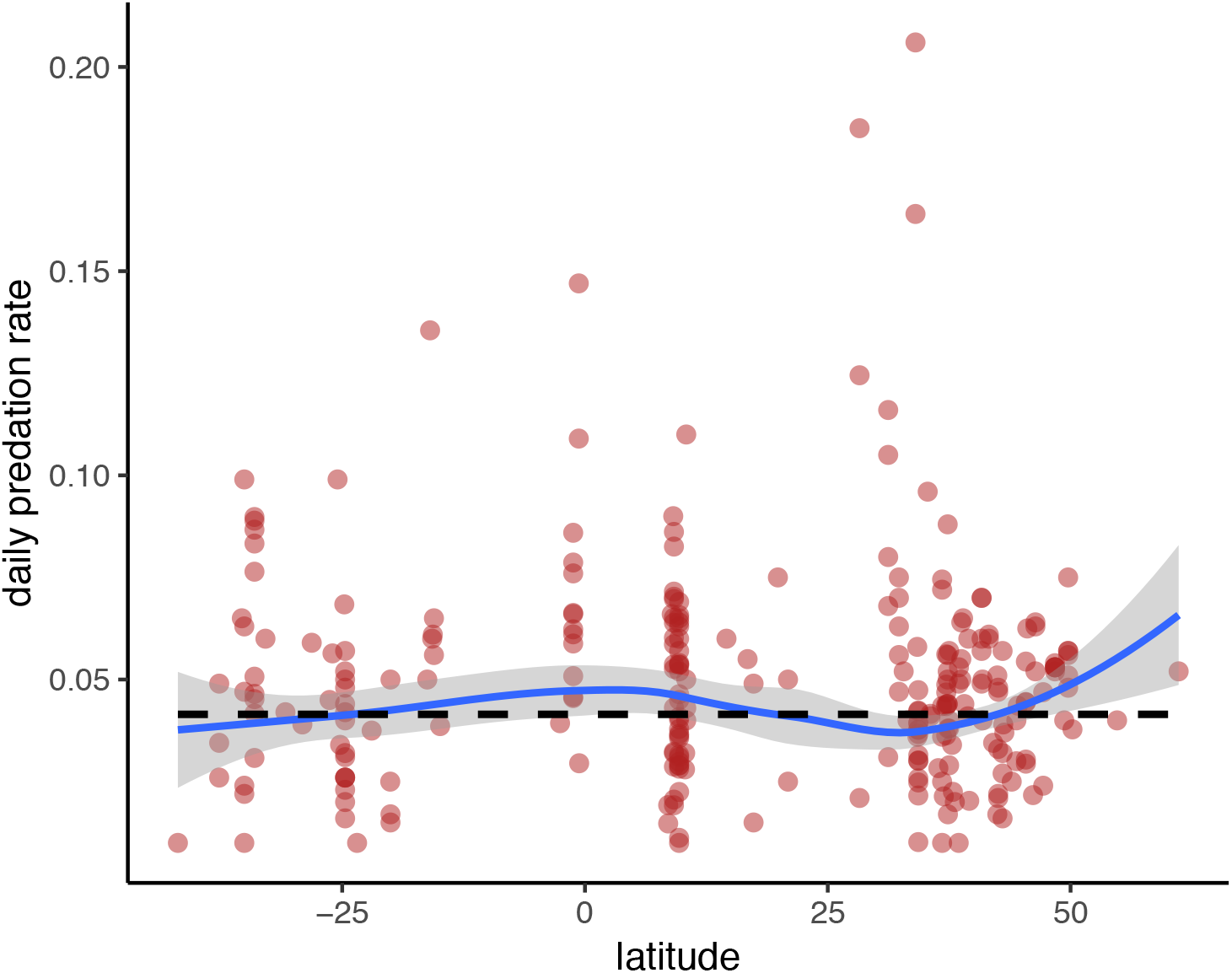
The latitudinal gradient in daily predation rates for land birds in the Americas (N = 267). Predictions from the best-fit metafor model are plotted as a dashed line—this simple model fit a constant value for daily predation rates across latitudes. For comparison, the loess fit, which used the same weights for each data point as the best-fit model, and incorporates the same estimated variance among the study-specific effect sizes, is plotted in blue with shaded 95% confidence intervals. Daily predation rates are estimated from daily survival rates reported in the literature.

### Nesting period length is a key covariate

We next conducted a series of exploratory analyses to investigate our finding of minimal latitudinal variation in daily predation rate. We first examined patterns in a conspicuous life history difference between latitudes, the length of the nesting period (Martin 2002; Chalfoun and Martin 2007). In our dataset, tropical nesting periods average approximately 15% longer than in the Northern temperate zone (~ 32 days vs ~ 28 days; Fig 3). This difference reflects a repeated pattern of evolution across a diversity of avian lineages. Latitude is negatively related to nesting period in a phylogenetic generalized least squares regression model (p < 0.0001, Table S3), and patterns are similar when analyzing only oscines (temperate origin, predominate in the temperate zone) or suboscines (tropical origin, predominate in the tropics; Table S6). An equal slopes model estimated that a 10-day increase in nesting period is associated with a 1.9% decrease in daily predation rates within each latitudinal zone (Table S4).

**Figure 3.**
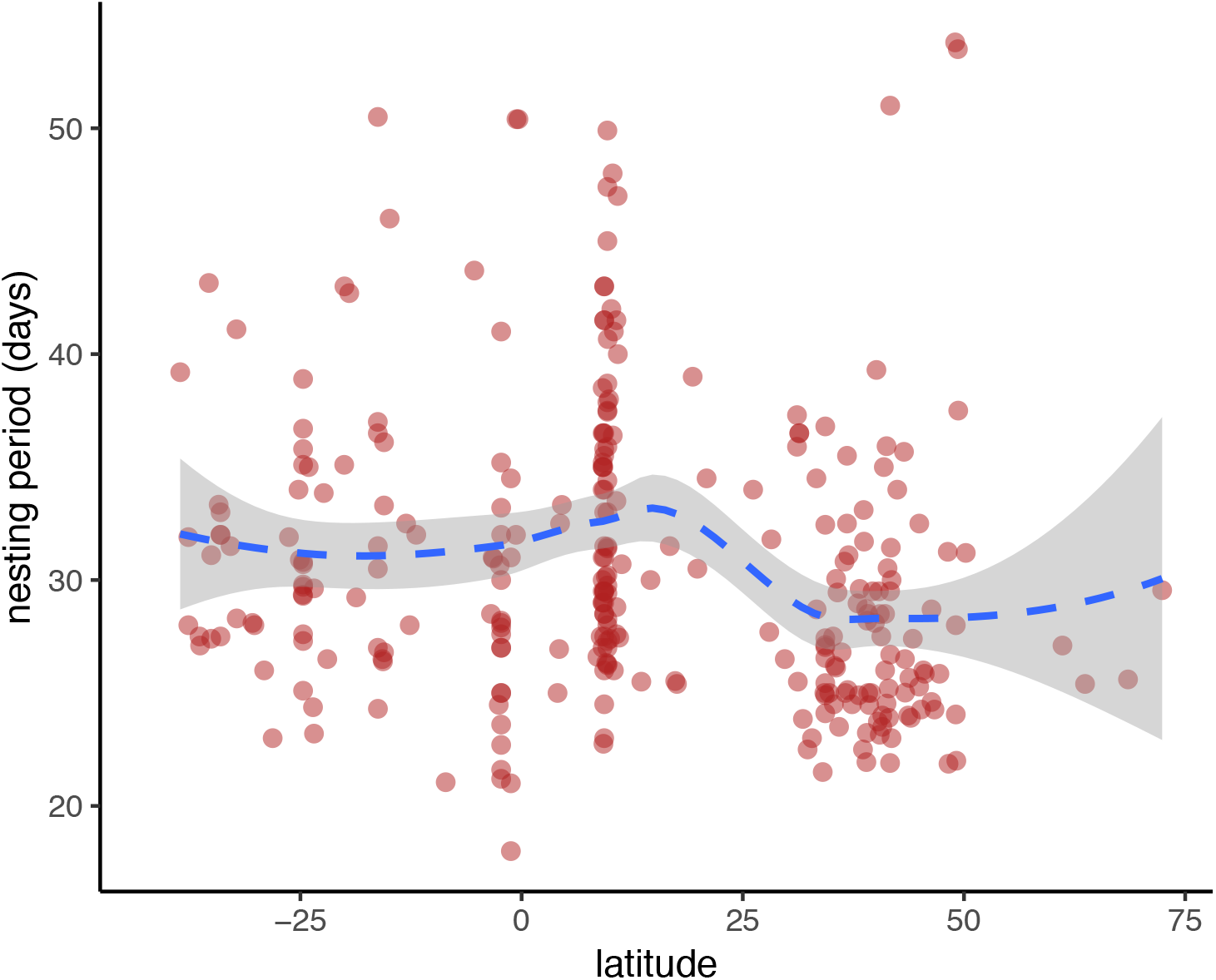
The latitudinal gradient in nesting period length for land birds in the Americas (N = 292 species). The dashed line illustrates the loess trendline, with 95% confidence intervals shaded in gray. Nesting periods average ~ 32 days within the tropics and Southern temperate zone but ~28 days within the Northern temperate zone. Nesting period is calculated as the number of days from the laying of the first egg to fledging.

We found that daily rates of nest predation are indeed greater in the tropics when we statistically controlled for length of nesting period (North temperate zone vs. tropics; p < 0.0001 in “same slopes” model to predict daily predation rate given latitudinal zone and nesting period; Fig 4, Table S4). This “same slopes” model was marginally better supported than a model that fit different slopes to different latitudinal zones (p = 0.092). When we controlled for nesting period length in the “different slopes” model, daily rates of nest predation were also greater in the tropics (comparing estimates for North temperate zone vs. tropics for the mean nesting period of 29 days, p < 0.001; Table S5, Fig S2). Again, this result is robust to potential sources of non-independence in our dataset (Table S6).

**Figure 4.**
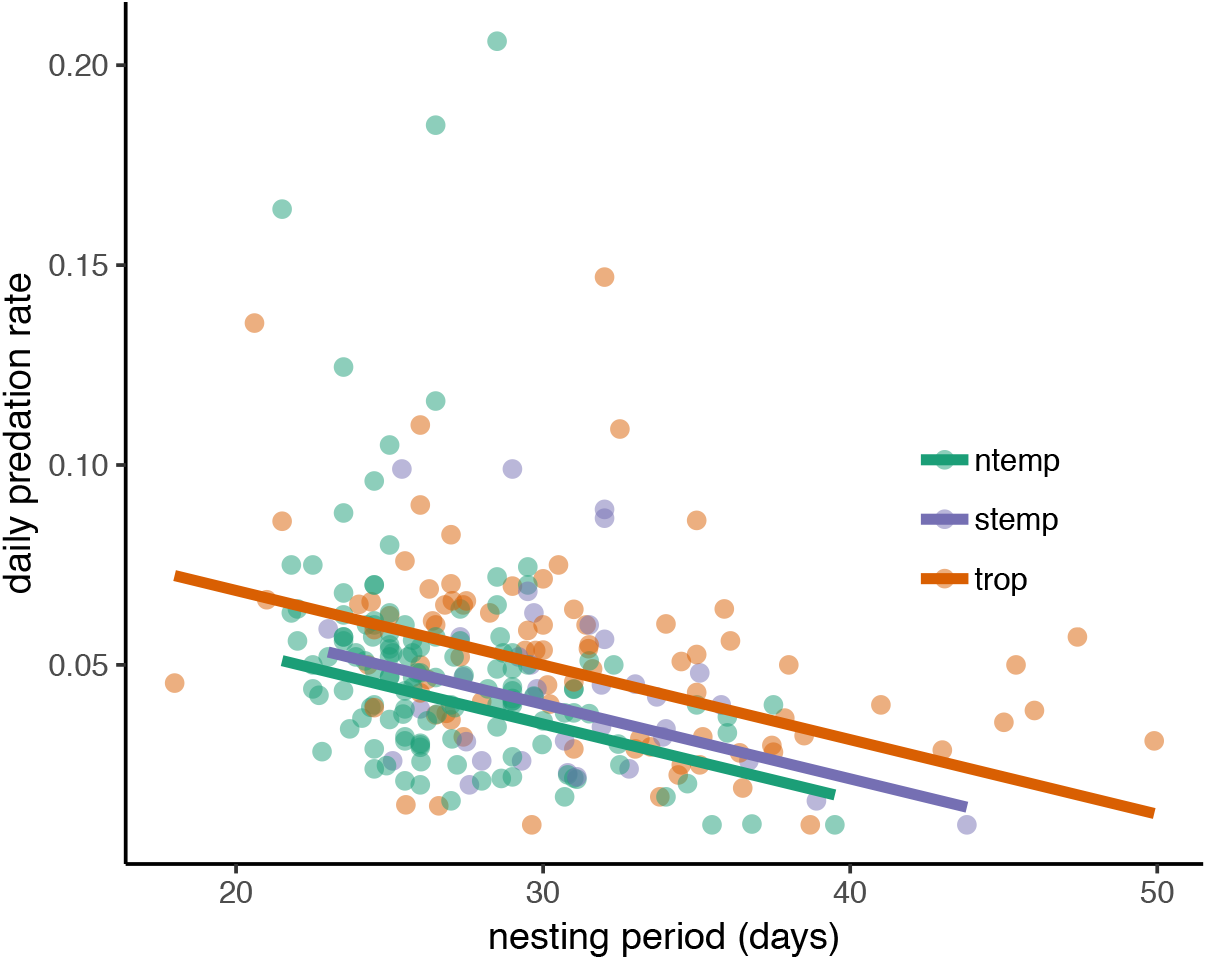
The relationship between daily predation rate and nesting period duration among latitudinal zones. Daily predation rate decreases with nesting period length in each latitudinal zone, but with different intercepts. Predictions from a metafor model are plotted as solid lines; values from individual studies are plotted in colors corresponding to their latitudinal zone (N = 252). Daily predation rates are estimated from daily survival rates reported in the literature; nesting period is calculated as the number of days from the laying of the first egg to fledging.

### Fledging success is lower in the tropics

The observation that nesting periods are longer in the tropics, while daily predation rates are similar across latitudes, implies that fledging success (the probability that a given nest survives to fledging) is lower in the tropics compared to the temperate zone. Indeed, we found strong evidence that fledging success of nests is highest in the Northern temperate zone (Fig 5, Table 2). A breakpoint regression estimated ~33% of nests successfully fledge nestlings within the tropics while ~ 50% of nests successfully fledge nestlings at 45° North (Table S7). Plotting this data revealed that this breakpoint regression model poorly fit data from the Southern temperate zone. A breakpoint regression model that fit different slopes to the Southern and Northern temperate zones was better supported (Table 2); this model estimated that ~34% of nests successfully fledge nestlings within both the tropics *and* the Southern temperate zone, while ~ 53% of nests successfully fledge nestlings at 45° N, a result that is robust to sources of non-independence in our data (Table S8). Although these differences in nest predation between the tropics and temperate zone seem consistent with the biotic interactions hypothesis, fledging success is integrated over a different nesting duration in the tropics than the temperate zone, and so is not an accurate measure of predation rate.

**Table 2.** Model comparison of metafor models with fledging success as the response variable. The best-fit model was a breakpoint regression that fit one slope within the tropics and unequal slopes in the Northern and Southern temperate zones.

| Model | $\Delta AIC$ |
| --- | --- |
| Breakpoint regression, unequal slopes (breakpoint = 23.4°) | -- |
| Breakpoint regression, equal slopes (breakpoint = 23.4°) | 4.68 |
| Linear regression | 21.29 |
| Constant (fledging success does not vary with latitude) | 66.35 |

**Figure 5.**
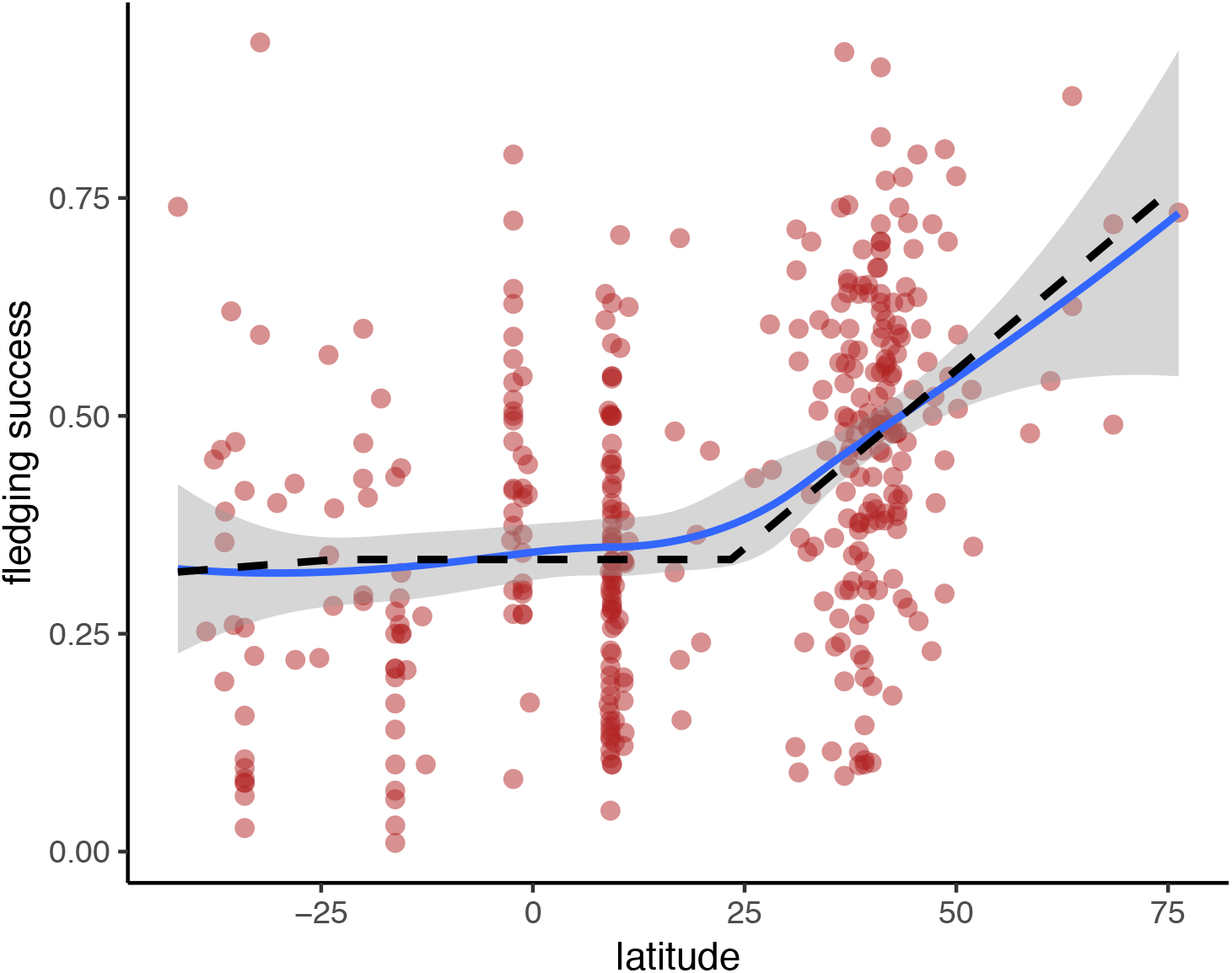
The latitudinal gradient in fledging success for land birds in the Americas (N = 367). Predictions from the best-fit model in metafor—a breakpoint regression that fit a flat line within the tropics and different slopes in the Northern and Southern Hemisphere temperate zones—, are plotted as a dashed line. For comparison, the loess fit, which used the same weights for each data point as the best-fit model, and incorporated the same estimated variance among the study-specific effect sizes, is plotted in blue with shaded 95% confidence intervals.

## Discussion

The “biotic interactions” hypothesis proposes that biotic interactions are stronger in the tropics, and that these strong interactions lead to faster evolution (and ultimately speciation) in the tropics. Inspired by these ideas, we ask whether adaptation may alter the link between ecological interaction and latitude. Our general hypothesis is that adaptation to different communities of interacting species at different latitudes can alter geographic patterns in interaction strength (see also Rasmann and Agrawal 2011; Chen et al. 2017). Hence, even if “baseline” interactions experienced by evolutionarily naïve populations are generally stronger in the tropics, evolved adaptations to different interaction regimes might flatten observed latitudinal gradients in interaction strength. We propose that this key role for adaptation may partially explain why some studies measuring interaction rates using “naïve” standardized models show higher interaction rates in the tropics (e.g., Roslin et al. 2017; Hargreaves et al. 2019) while studies on wild “adapted” populations often show minimal variation between interaction rate and latitude (reviewed by Moles and Ollerton 2016).

Our data are consistent with a potential role for adaptation in influencing latitudinal patterns in interaction rates. We report that there is little or no latitudinal gradient in daily rates of nest predation. At the same time, there is a latitudinal gradient in daily rates of nest predation when statistically controlling for nesting period duration. Reconciling these results requires explicit consideration of the fact that birds at different latitudes possess different life history adaptations that mitigate baseline differences in nest predation rates. Tropical birds have longer nesting periods, and longer nesting periods are generally associated with lower daily predation rates. The implication is that if birds at all latitudes had the same nesting period length, then tropical birds would suffer higher rates of nest predation than temperate zone birds. We view this scenario—where we statistically control for latitudinal divergence in a life history trait—as a measurement of “baseline” interaction strength. However, in reality, birds are not equivalent across latitudes. Instead, tropical and temperate species differ in life history adaptations including nesting period length. When taking consideration of different adaptations along the gradient, tropical birds no longer suffer higher rates of nest predation, and hence the latitudinal gradient in “derived” interaction strength is more flat. A prediction arising from this interpretation is that a standardized study of predation on artificial nests (which are “evolutionarily naïve”) across latitudes would find higher rates of nest predation in the tropics. The only such standardized study we are aware of was conducted at multiple latitudes within the Arctic. Consistent with our expectation, this study reported higher predation rates on artificial nests at lower latitudes (Mckinnon et al. 2010).

Our interpretation of latitudinal patterns in nest predation assumes that longer nesting periods in tropical birds represent in part an adaptation to tropical predator assemblages. Substantial evidence suggests that predation can be an important driver of latitudinal differences in avian life history traits (Ghalambor and Martin 2001; Martin 2002; Martin 2015). Tropical predator assemblages could select for long nesting periods in tropical birds relative to temperate birds either directly or indirectly. For example, high baseline predation risk in the tropics could directly select for reduced rates of parental care. Less parental care per day at the nest is often associated with longer nesting periods (Chalfoun and Martin 2007; Martin et al. 2007; but see Tieleman et al. 2004), and the tendency for birds to reduce their rates of activity at the nest when confronted with high predation risk is especially strong in the tropics (Matysioková and Remeš 2018). Perhaps more plausibly, tropical predator assemblages could indirectly select for long nesting periods in tropical birds via its correlation with a suite of life history traits. For example, high predation could directly select for increased investment in adult survival, with longer nesting periods as an indirect effect (e.g., by a trade-off between adult investment in survival vs. incubation/provision rates; Martin et al. 2015). The exact mechanism by which predator-driven selection (among many sources of selection on life history traits) favors longer nesting periods in tropical birds is not crucial to our analysis. Our general idea is that adaptation to gradients in baseline interaction regimes can alter observed rates of interaction. Applied to the bird nest predation data we present here, our assumption is simply that variation in nesting period duration across latitudes represents a component of adaptation to changing predator communities.

### Comparison with previous studies of nest predation across latitudes

We report that natural rates of nest predation experienced by land birds are similar along a latitudinal gradient in the Americas stretching from ~ 40**°** South to ~ 60**°** North. This finding surprised us, because nest predation has been considered an archetypical example in support of the more general hypothesis that biotic interactions are stronger at low latitudes (Schemske 2009). Here we show that, at least for Western Hemisphere land birds, the long-held view that nest predation is higher in the tropics appears to be derived from observations of reduced fledging success of tropical birds. However, latitudinal differences in fledging success are a consequence of evolved differences between birds at different latitudes in a life history trait (nesting period) rather than of differences in interaction rates. We focus on daily predation rates because analyzing latitudinal patterns in a biotic interaction requires a common currency for measuring interaction strength (Bulla et al. 2018). Whether the pattern we document for the Western Hemisphere applies to the Eastern Hemisphere remains uncertain; for example, rates of nest predation are indeed higher in the tropics in Australia (Remeš et al. 2012; note that this study found much greater longitudinal variation in nest predation than latitudinal variation).

### Conclusions

The biotic interactions hypothesis invokes strong biotic interactions in the tropics to explain in part the high species richness of the tropics. However, whether interactions are stronger in the tropics remains hotly debated. On one hand, one review reported general support for stronger biotic interactions in the tropics (Schemske et al. 2009). On the other hand, a different review suggested the notion that interactions are stronger in the tropics to be a “zombie idea”—an idea that is not supported by empirical data but that refuses to die (Moles and Ollerton 2016). We propose that adaptation that alters baseline interaction rates may in part reconcile these opposing viewpoints (note that methodological differences are another factor responsible for variation in the outcomes of observational studies). From this perspective, we expect interactions to be stronger in the tropics when studies use standardized evolutionarily naïve models to measure it (e.g., clay caterpillars or sunflower seeds; Roslin et al. 2017; Hargreaves et al. 2019), but not necessarily when studies measure interaction strengths experienced by wild evolutionarily “adapted” populations.

More broadly, we argue that documenting geographic patterns in interaction strength is only part of the story. The biotic interactions hypothesis is intriguing precisely because it links biotic interactions to rates of divergence to explain geographic patterns in species richness. We propose that variation in biotic interactions across latitudes might exert strong selection for differential adaptations along the gradient, and that these adaptations may reduce or flatten latitudinal gradients in interaction strength. If so, adaptations could potentially flatten any latitudinal gradient in evolutionary rates as well, though this remains untested. We advocate for an increased focus on the evolutionary consequences of variation in the strength of biotic interactions (Benkman 2013). Moving forward, we suggest that the most rigorous tests of the biotic interaction hypothesis will consider how both biotic interaction strength and evolutionary rates of trait divergence vary across latitude.

## Acknowledgements and Funding

This research was supported by postdoctoral fellowships from the Biodiversity Research Centre and Banting Canada (#379958) to BGF. None of our funders had any influence on the content of the submitted manuscript, and none of our funders required approval of the final manuscript to be published. Comments from the Schluter lab group and Ralf Yorque greatly improved this manuscript.

---

**Figure S1.**
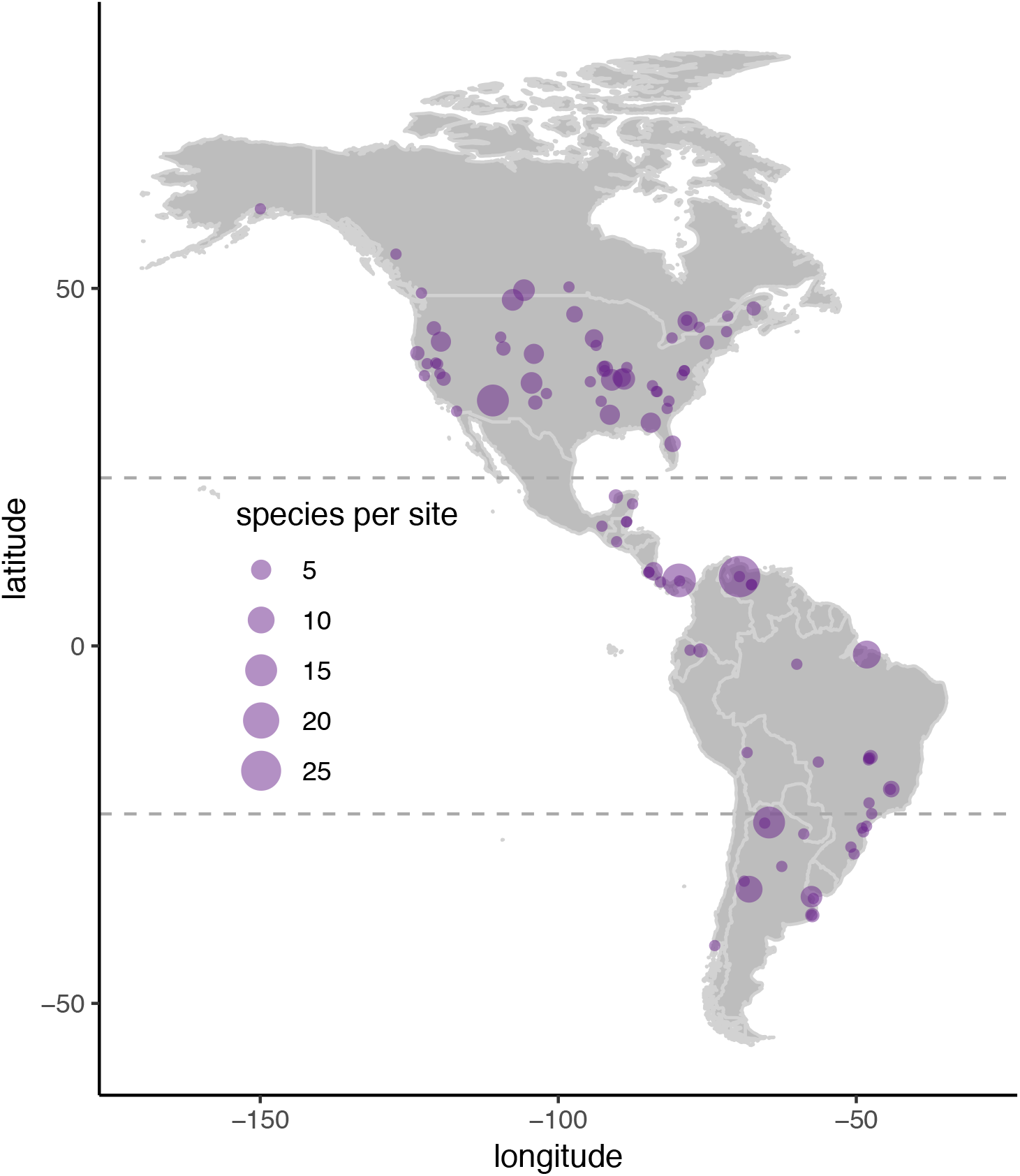
Map of studies measuring daily rates of nest survival for land birds in the Americas (dataset = 110 studies). We used these data to estimates daily rates of nest predation, as the bulk of nest failure across latitudes is due to predation. Many studies report data for multiple species from the same site, illustrated by the size of the circle. The Tropics of Cancer and Capricorn (at 23.4**°** N and S, respectively) delimit the tropics, and are illustrated with dashed lines.

**Figure S2.**
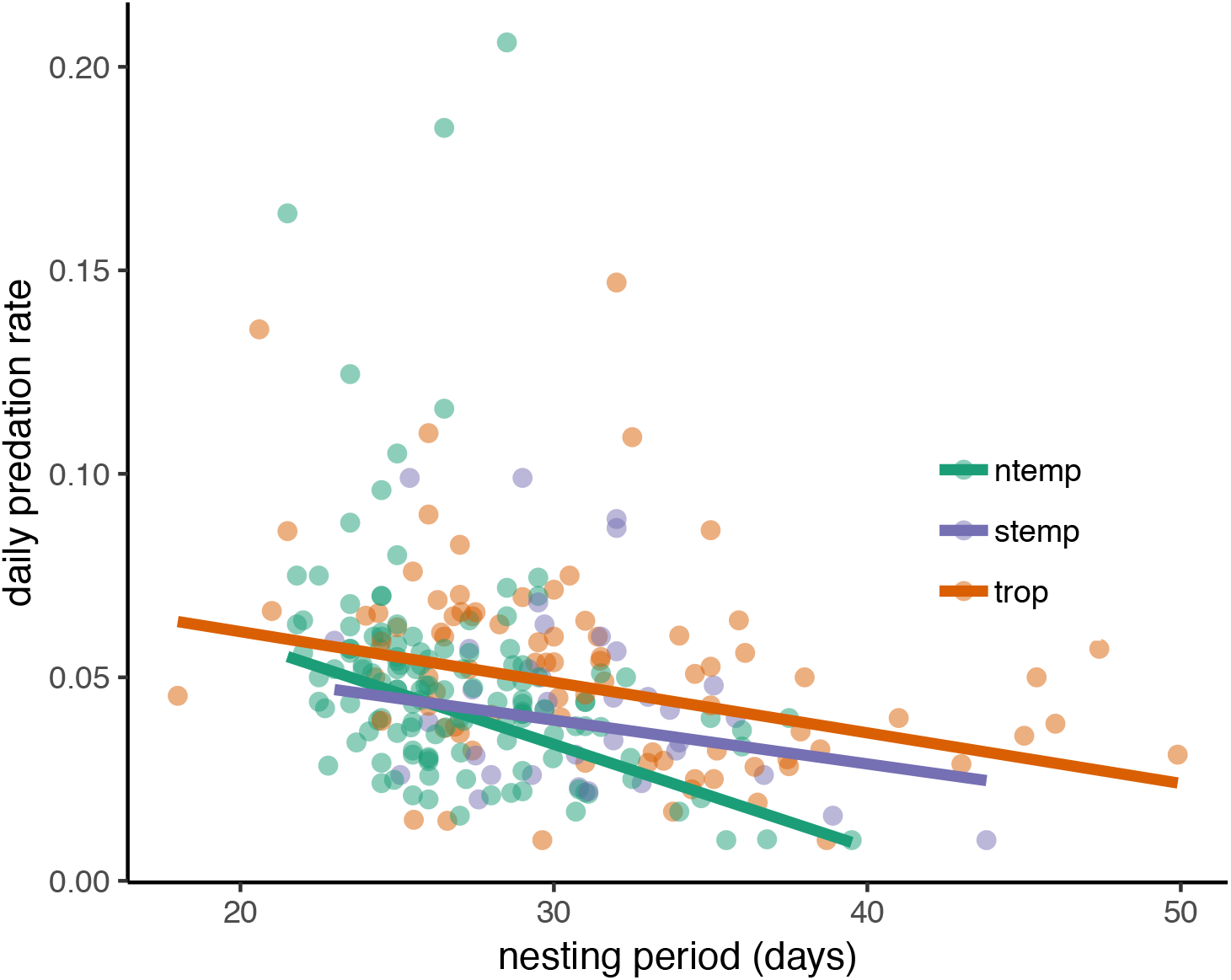
The relationship between daily predation rate and nesting period duration among latitudinal zones (different slopes model). Daily predation rate decreases with nesting period length, but with different intercepts and slopes for different latitudinal zones. Predictions from a metafor model are plotted as solid lines; values from individual studies are plotted in colors corresponding to their latitudinal zone (N = 252). Daily predation rates are estimated from daily survival rates reported in the literature; nesting period is calculated as the number of days from the laying of the first egg to fledging.

**Table S1.** Parameter estimates for fixed effects from metafor models with daily predation rate as the response variable. Slope estimates for the linear regression are for straight lines outward from the equator towards higher latitudes; slope estimates for breakpoint regression are for straight lines outward from the tropics (23.4 degrees latitude) towards higher latitudes.

| Model | Intercept ± SE | Slope ± SE |
| --- | --- | --- |
| Intercept-only<br>(constant) | 0.042 ± 0.0015 | NA |
| Linear regression | 0.044 ± 0.0039 | -0.0001 ± 0.0001 |
| Breakpoint<br>regression<br>(breakpoint = 23.4°) | 0.041 ± 0.0024 | 0.0001 ± 0.0002 |

**Table S2.** Model summary of intercept-only metafor model with daily predation rate as the response variable and species and phylogeny as random effects.

| Effect | Estimate |
| --- | --- |
| Intercept (fixed) | 0.042 (se = 0.0083) |
| Species (random) | 0.0000 (sqrt = 0.0067) |
| Phylogeny (random) | 0.0003 (sqrt = 0.016) |

**Table S3.** Parameter estimates for fixed effects from phylogenetic generalized least squares regression models with nesting period as the response variable. Pagel’s λ for these model was 0.99 (entire dataset), 1.00 (oscines) and 0.94 (suboscines).

| Model | Parameter | Estimate ± SE |
| --- | --- | --- |
| Entire dataset | Intercept | 27.29 ± 2.90 |
| | Absolute value (latitude) | -0.040 ± 0.015 |
| Oscines only | Intercept | 34.92 ± 2.86 |
| | Absolute value (latitude) | -0.026 ± 0.016 |
| Suboscines only | Intercept | 35.45 ± 2.99 |
| | Absolute value (latitude) | -0.065 ± 0.039 |

**Table S4.**
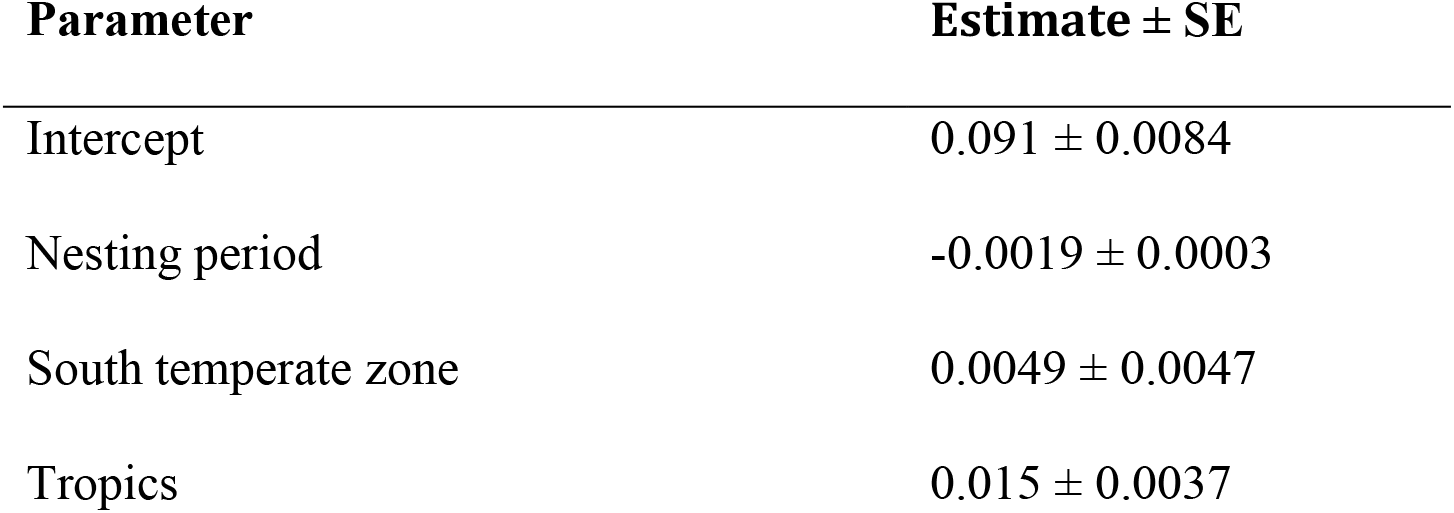
Parameter estimates from a metafor model with daily predation rate as the response variable. This is the “equal slopes” model where different latitudinal zones have the same slope. The reference category is the North temperate zone; parameter estimates for the South temperate zone and tropics represent deviations from this reference category.

| Parameter | Estimate ± SE |
| --- | --- |
| Intercept | 0.091 ± 0.0084 |
| Nesting period | -0.0019 ± 0.0003 |
| South temperate zone | 0.0049 ± 0.0047 |
| Tropics | 0.015 ± 0.0037 |

**Table S5.** Parameter estimates from a metafor model with daily predation rate as the response variable. This is the “different slopes” model where different latitudinal zones have different slopes. The reference category is the North temperate zone; parameter estimates for “Nesting Period: South temperate zone” and “Nesting Period: Tropics” represent deviations from this reference category.

| Parameter | Estimate ± SE |
| --- | --- |
| Intercept | 0.10 ± 0.012 |
| Nesting period | -0.0025 ± 0.00043 |
| South temperate zone | -0.038 ± 0.035 |
| Tropics | -0.024 ± 0.019 |
| Nesting period: South temperate zone | 0.0015 ± 0.0012 |
| Nesting period: Tropics | 0.0013 ± 0.00063 |

**Table S6.** Parameter estimates from a metafor model with daily predation rate as the response variable and study, species and phylogeny as random effects. This is the “equal slopes” model where different latitudinal zones have the same slope. The reference category for the intercept is the North temperate zone; parameter estimates for the South temperate zone and tropics represent deviations from this reference category.

| Parameter | Estimate ± SE |
| --- | --- |
| Intercept | 0.091 ± 0.0084 |
| Nesting period | -0.0019 ± 0.00030 |
| South temperate zone | 0.0050 ± 0.0047 |
| Tropics | 0.015 ± 0.0037 |
| Species (random) | 0.0 (sqrt = 0.0064) |
| Phylogeny (random) | 0.0 (sqrt = 0.0) |

**Table S7.**
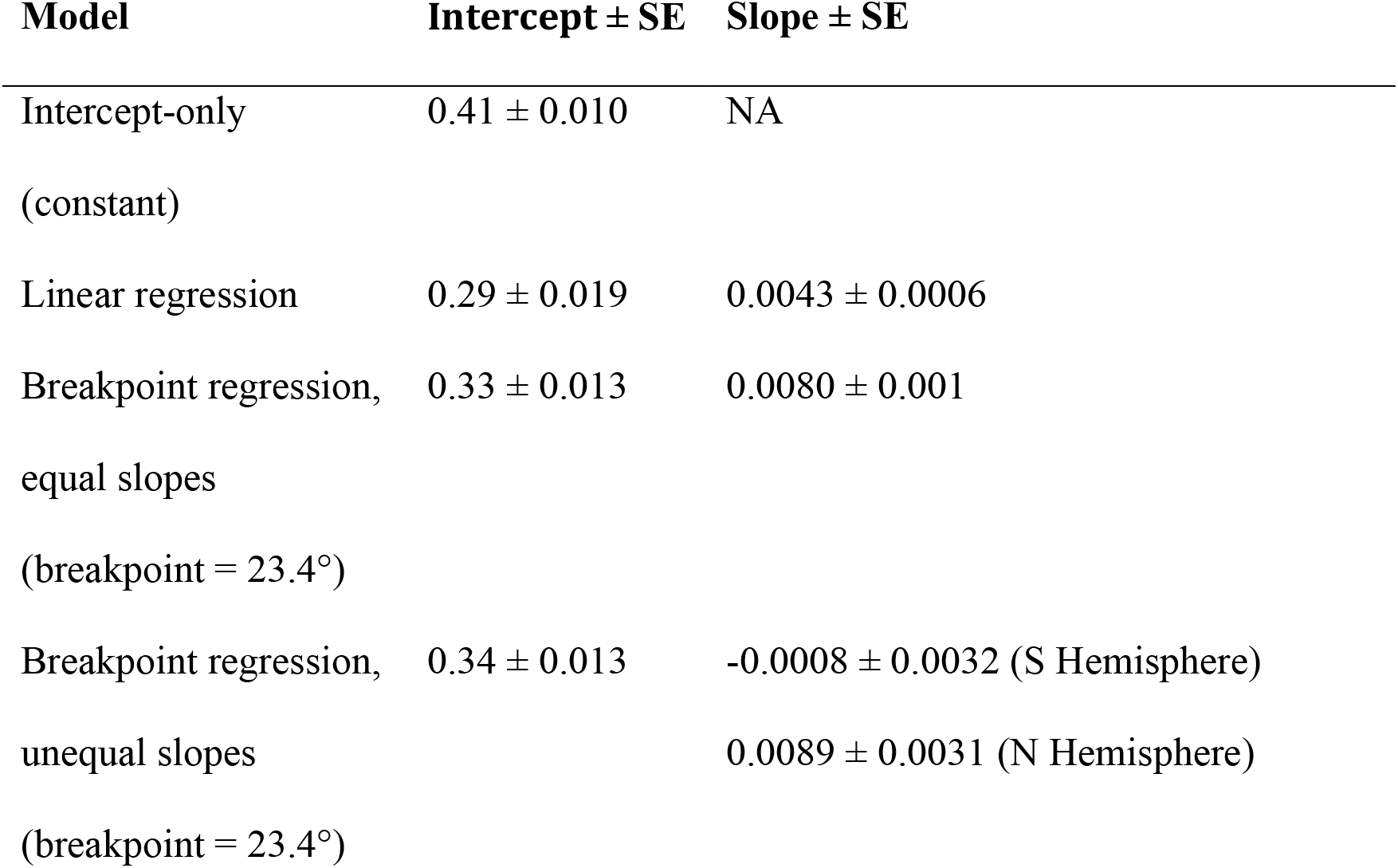
Parameter estimates for fixed effects from metafor models with fledging success as the response variable. Slope estimates for the linear regression are for straight lines outward from the equator towards higher latitudes; slope estimates for breakpoint regression are for straight lines outward from the tropics (23.4 degrees latitude) towards higher latitudes. Breakpoint regressions fit either equal slopes (same slope value in both Northern and Southern temperate zones) or unequal slopes (different slopes in Northern vs. Southern temperate zones).

**Table S8.** Model summary of unequal slopes breakpoint regression metafor model with fledging success as the response variable, and species and phylogeny included as random effects.

| Effect | Estimate |
| --- | --- |
| Intercept (fixed) | 0.34 (se = 0.013) |
| Slope (fixed) | -0.0008 ± 0.0032 (S Hemisphere) |
|  | 0.0089 ± 0.0031 (N Hemisphere) |
| Species (random) | 0.026 (sqrt = 0.16) |
| Phylogeny (random) | 0 (sqrt = 0.0000) |

